# Molecular Evolution of the Sex Peptide Network in *Drosophila*

**DOI:** 10.1101/833939

**Authors:** Meaghan K. McGeary, Geoffrey D. Findlay

## Abstract

Successful reproduction depends on interactions between numerous proteins beyond those involved directly in gamete fusion. While such reproductive proteins evolve in response to sexual selection pressures, how networks of interacting proteins arise and evolve as reproductive phenotypes change remains an open question. Here, we investigated the molecular evolution of the “sex peptide network” of *Drosophila melanogaster*, a functionally well-characterized reproductive protein network. In this species, the peptide hormone sex peptide (SP) and its interacting proteins cause major changes in female physiology and behavior after mating. In contrast, females of more distantly related *Drosophila* species do not respond to SP. In spite of these phenotypic differences, we detected orthologs of all network proteins across 22 diverse *Drosophila* species and found evidence that most orthologs likely function in reproduction throughout the genus. Within SP-responsive species, we detected the recurrent, adaptive evolution of several network proteins, consistent with sexual selection acting to continually refine network function. We also found some evidence for adaptive evolution of several proteins along two specific phylogenetic lineages that correspond with increased expression of the SP receptor in female reproductive tracts or increased sperm length, respectively. Finally, we used gene expression profiling to examine the likely degree of functional conservation of the paralogs of an SP network protein that arose via gene duplication. Our results suggest a dynamic history for the SP network in which network members arose before the onset of robust SP-mediated responses and then were shaped by both purifying and positive selection.

## Introduction

Successful reproduction requires the fusion of egg and sperm cells, yet this fusion is often facilitated by proteins that are not part of the gametes. For example, non-gametic reproductive proteins provided in male seminal fluid or produced in the female reproductive tract can facilitate sperm motility, induce or manage sperm storage or cause changes to female reproductive physiology (Wilburn & Swanson, 2016, Schnakenberg et al., 2011). While proteomic and comparative genomic methods have enabled the identification of hundreds of gametic and non-gametic reproductive proteins across diverse taxa (reviewed in McDonough et al., 2016), understanding how these proteins interact, and how such interactions evolve, remain areas of active research.

Some of the best-characterized reproductive protein interactions occur in the “sex peptide network” of *Drosophila melanogaster* that regulates female post-mating behavior and physiology. The network centers on the sex peptide (SP), a short peptide hormone transferred from males to females as a non-gametic component of seminal fluid (Chen et al., 1988). The presence of SP in the female reproductive tract stimulates egg production (Soller et al., 1999), reduces receptivity to remating (Liu & Kubli, 2003, Chapman et al., 2003), facilitates the release of sperm from storage prior to fertilization (Avila et al., 2010), and affects numerous other female behaviors, including feeding, defecation and sleep (Carvalho et al., 2006, Apger-McGlaughon & Wolfner, 2013, Isaac et al., 2010). SP-mediated effects on females persist for several days after mating because SP binds to sperm, which become stored in specialized storage organs in the female tract (Peng et al., 2005). SP is then gradually cleaved from sperm and released from the storage organs into the female tract, where it interacts with the sex peptide receptor (SPR), a G-protein coupled receptor that is expressed in a subset of neurons innervating the uterus (Yapici et al., 2008, Hasemeyer et al., 2009, Yang et al., 2009). This gradual “dosing” of SP causes the persistence of the hormone’s effects on female behavior and physiology. SPR signaling is also required for the efficient release of sperm from the storage organs (Avila et al., 2015).

While the molecule(s) on sperm to which SP binds remain unknown, RNAi screens have identified several additional male seminal fluid proteins and female reproductive tract proteins required for robust SP responses (Ravi Ram & Wolfner, 2007, Ravi Ram & Wolfner, 2009, LaFlamme et al., 2012, Findlay et al., 2014, Singh et al., 2018). Together with SP and SPR, these proteins comprise the SP network. The male-derived proteins include: predicted C-type lectins CG1652 and CG1656; predicted proteases/protease homologs CG9997, seminase, aquarius and intrepid; and, predicted cysteine-rich secretory proteins CG17575 and antares. The female-derived proteins include fra mauro (a predicted metallopeptidase), Esp (a predicted anion transporter) and hadley (which lacks identifiable protein domains). The male-derived proteins act interdependently to facilitate SP binding to sperm (Ravi Ram & Wolfner, 2009, Findlay et al., 2014, Singh et al., 2018), while the female-derived proteins act downstream of SP binding to sperm, potentially by facilitating SP-SPR signaling (Findlay et al., 2014). Other genes expressed in the secondary cells of the male accessory gland are also required for SP-mediated responses, though it remains unclear whether these genes encode proteins that interact directly with the network proteins described above (Sitnik et al., 2016).

SP’s functions and interactions have been well characterized in *D. melanogaster*, but comparative genomic and functional studies have shown that the SP response is not conserved throughout the *Drosophila* genus. Tsuda et al. (2015) found that only species of the *melanogaster* group of *Drosophila* (Fig. 1) show changes in female remating receptivity and egg production upon injection with synthetic SP, even though SP and SPR orthologs can be detected outside of this group. Furthermore, by incubating GFP-labeled SP with female reproductive tracts from progressively more divergent species, Tsuda et al. (2015) discovered that SP could bind to the female tract only in *melanogaster* group species. This observation suggested that robust expression of SPR in the female tract evolved on the phylogenetic lineage leading to the *melanogaster* group, which the authors tested by comparing *SPR* gene expression between in-group and out-group species (Tsuda et al., 2015). Consistent with *D. melanogaster* expression patterns (Yapici et al., 2008), they found that *SPR* was expressed in non-reproductive areas in both sexes of all species examined. However, its expression in the female reproductive tract was largely limited to the *melanogaster* group. (The only outgroup species that showed expression in this location was *D. virilis*, but conspecific GFP-labeled SP did not bind to female reproductive tracts in this species.) Intriguingly, the SP ortholog from *D. pseudoobscura* (a non-*melanogaster* group species) is expressed in *D. pseudoobscura* male reproductive tracts (Yang et al., 2018) and can trigger SP-mediated responses when injected into *D. melanogaster* females, but not when injected into conspecifics (Tsuda et al., 2015). This result suggests that the SP protein might have evolved the potential to affect female post-mating behavior before the emergence of the *melanogaster* group, but this function was not fully realized until the subsequent evolution of SPR expression in the female reproductive tract (and, perhaps, within specific neurons in the tract) (Hasemeyer et al., 2009, Yang et al., 2009, Yapici et al., 2008, Rezaval et al., 2012). It is also possible that the transition to high levels of SPR expression in the female reproductive tract created or intensified an evolutionary selective pressure to bind higher levels of SP to stored sperm.

**Figure 1.**
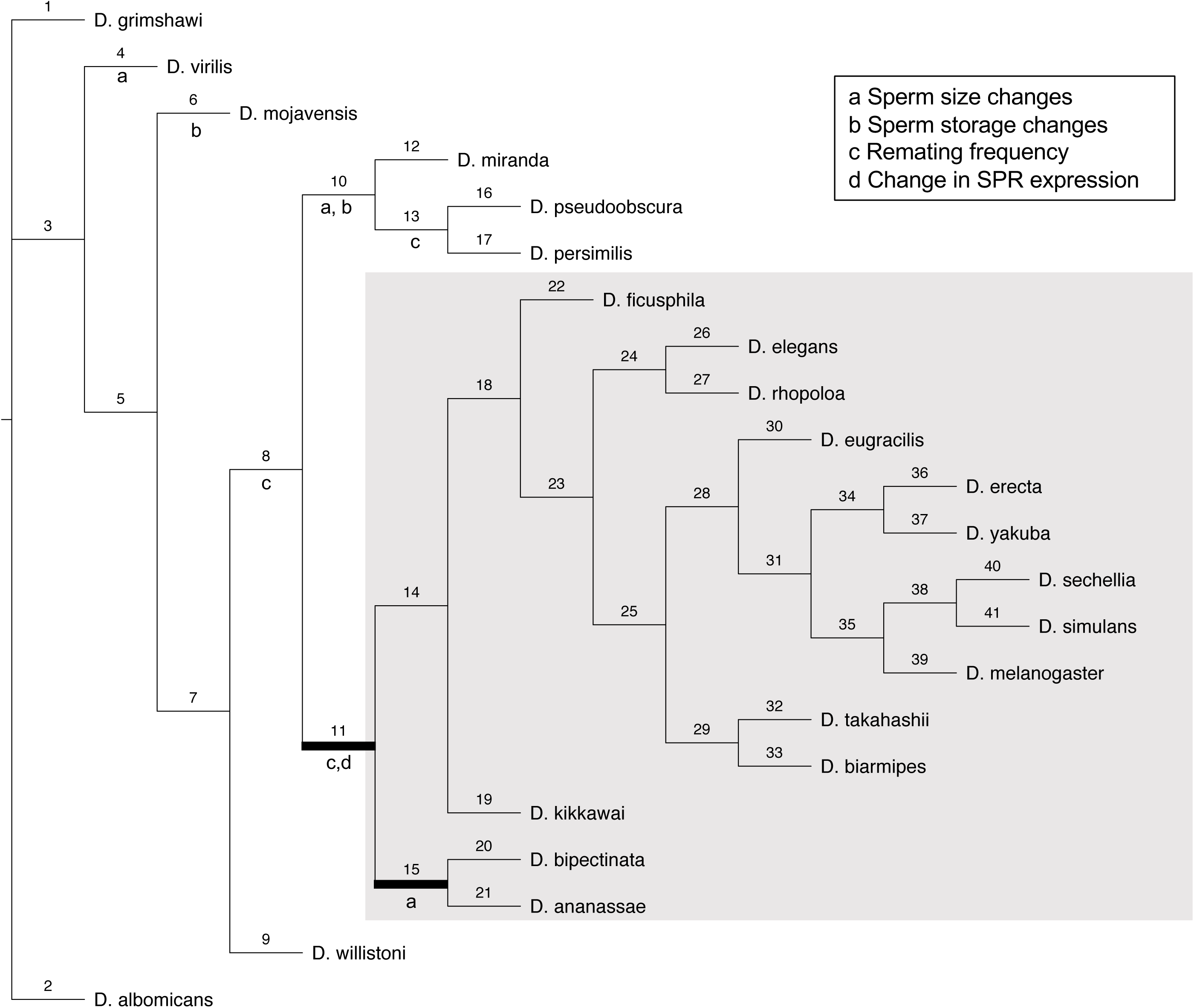
Phylogeny of *Drosophila* species examined in this study. The gray box indicates the *melanogaster* group. Each branch is numbered for reference in the main text. Key changes in reproductive tracts are indicated by letters a-d and are based on an examination of the literature (references cited in main text). The PAML branch-sites tests (see Results) were conducted on branches 11 and 15. Branch lengths are not proportional to evolutionary distances.

While *Drosophila* species differ in reproductive traits for many reasons (Markow & O’Grady, 2005), some of these differences relate directly to the SP network and could thus be causes or consequences of SP network evolution. For example, a change in sperm length may affect the amount of SP that can bind, and other structural changes to sperm could affect the binding of SP and other network proteins that interact with sperm, such as CG1652 and CG1656 (Ravi Ram & Wolfner, 2009, Singh et al., 2018). The amount of SP bound to sperm, the rate and efficacy of its release (Peng et al., 2005), and its ability to bind SPR (Yapici et al., 2008) could affect re-mating rates, while the structure of female sperm storage organs could affect the ability of the network proteins to bind SP to sperm or modulate SP’s interaction with SPR. Changes in these traits – sperm length, female remating rate, and female sperm storage structures – have been well documented in the literature (Alpern et al., 2019, Joly & Bressac, 1994, Markow, 1996, Markow & O’Grady, 2005, Pitnick et al., 1999, Snook et al., 1994, Snook, 1995, Singh et al., 2002), and we summarize them and infer their timing in Figure 1. Of particular relevance to this study, the phenotypic and phylogenetic data are consistent with SPR evolving to be expressed in female reproductive tracts along the lineage leading to the *melanogaster* group of flies (Tsuda et al., 2015; branch 11) and an increase in sperm length on the lineage leading to *D. ananassae* and *D. bipectinata* (Joly & Bressac, 1994, Markow, 1996, Pitnick et al., 1999; branch 15).

In light of the differences between species in reproductive phenotypes, we used comparative genomics and molecular evolutionary analysis to gain insights into the evolution of the SP network. While robust, long-lasting changes in female behavior and physiology due to SP are found only in the *melanogaster* group of *Drosophila*, we identified orthologs of each SP network protein in numerous outgroup species and verified their expression in the male reproductive system in two such species from published data (Yang et al., 2018, Kelleher et al., 2009). Using PAML (Yang, 2007) we determined that recurrent positive selection has acted on specific sites in several of these proteins. We also detected marginal evidence that positive selection has acted on certain network proteins on key phylogenetic lineages corresponding with major changes in SP-related phenotypes. Finally, we used RT-PCR to investigate whether any gross-scale changes in gene expression occurred surrounding the gene duplication event that gave rise to one of the SP network proteins, seminase. Taken together, our results suggest that the members of the SP network had the potential to influence reproductive success before the onset of SP/SPR-mediated responses in the reproductive tract of mated females of the *melanogaster* group of species. However, additional adaptive changes in these proteins occurred concurrent with, and subsequent to, these critical changes in the fly reproductive system. These results underscore the strength of sexual selection acting in *Drosophila* and illustrate potential molecular changes that occur in the face of such selection.

## Methods

### Identification of SP network proteins across Drosophila species

We obtained the protein sequence for each SP network protein in *D. melanogaster* from FlyBase. For species for which protein annotations were available on FlyBase (Drosophila 12 Genomes Consortium, 2007), we obtained orthologous protein-coding DNA sequences using the FlyBase Orthologs feature. These species included *Drosophila simulans, sechellia, yakuba, erecta, ananassae, pseudoobscura, persimillis, willistoni, mojavensis, virilis* and *grimshawi.* For species with sequenced genomes that lacked FlyBase protein annotations (Chen et al., 2014), we manually searched for gene orthologs using tBLASTn and the *D. melanogaster* protein sequence as the query. These species include *Drosophila ficusphila, eugracilis, takahashii, elegans, rhopoloa, kikkawai, bipectinata, miranda* and *albomicans.* For genes expected to have introns based on the *D. melanogaster* gene structure, we looked in the unannotated species for the approximate location of the *D. melanogaster* intron, and used known intron border consensus sequences and six-frame translation, implemented in EMBOSS SixPack (Madeira et al., 2019), to identify predicted intron borders and remove intronic sequences prior to the analyses below.

To study the gene duplication events that gave rise to *seminase* and its tandem gene duplicates (*CG10587* and *CG11037* in *D. melanogaster*), we identified the genes flanking these three genes and used them to identify the syntenic region of the other *Drosophila* genomes. We assumed conservation of gene order within this syntenic region in assigning orthologs for this gene family (Figure S1).

For all putative orthologs identified by bioinformatic methods, we verified that the ortholog was the reciprocal best BLAST hit to the expected SP network member of *D. melanogaster.* Inferred orthologs with a high degree of similarity, successful reciprocal best hits, and a sequence that could be translated conceptually to produce a polypeptide without premature stops, were retained for study. In cases of duplicate genes (*seminase*, *CG1652* and *CG1656*), we also used gene order and synteny to confirm correct ortholog identification.

### Sequence alignment

For each SP network protein, we used MUSCLE as implemented in MEGA 6.06 (Tamura et al., 2013) to align amino acid sequences, then visually checked and edited each alignment for accuracy. Amino acid alignments were then back-translated in MEGA to obtain the cDNA alignment.

### Phylogenetic analysis

To infer a *Drosophila* consensus phylogeny based on all SP network proteins, we concatenated the amino acid alignments of all SP network proteins within each of the 22 species. We used PROML in Phylip (Felsenstein, 2005) to infer an unrooted maximum-likelihood phylogeny (with random input order, slow analysis, and all other default parameters). Gaps in the alignment were used in cases in which a protein was not present in a particular species. The resulting phylogeny matched published *Drosophila* phylogenies, except for *D. virilis* and *D. mojavensis* (Drosophila 12 Genomes et al., 2007, Markow & O’Grady, 2005, Seetharam & Stuart, 2013). We then used this consensus tree for the PAML analyses, with species removed on a gene-by-gene basis as described below.

### Detection of recombination

Because recombination within a gene sequence can impact the results of analyses to detect selection, we first used GARD with default parameters as implemented in DataMonkey 2.0 to check for evidence of recombination within each gene (Kosakovsky Pond et al., 2006, Weaver et al., 2018). Genes were partitioned at breakpoints evaluated as significant by the Kishino-Hasegawa test (p-value < 0.05 for both LH and RH; Table S2), and PAML was run on each gene segment separately. We performed PAML analyses on sequence alignments spanning two different ranges in the *Drosophila* phylogeny: the branch and branch-sites tests (see below) were run on species from the entire genus, while the sites test (see below) was run on species from only the *melanogaster* group. Thus, we generated a set of recombination breakpoints for each set of species (i.e., all examined members of the genus or members of the *melanogaster* group; Table S2). For each species set, six SP network genes showed evidence of recombination, but the genes that showed recombination differed between the two sets of species.

### PAML analyses

For each protein, we used codeml of the PAML package to perform evolutionary analyses on protein-coding DNA sequence (Yang, 2007). To test for heterogeneity in the rate of each protein’s evolution across the phylogeny, we utilized the PAML branch test, which uses a likelihood ratio test (LRT) to compare the “free ratio” model, allowing for different ω values for each branch, with model M0, which estimates a single ω for the whole phylogeny (Yang, 1998). For these tests, and for the branch-sites tests below, we used the consensus tree described above that covered the entire *Drosophila* phylogeny, but manually removed from it any species for which: a) an ortholog could not be identified, or b) an ortholog was identified, but it could not be confidently aligned due to ambiguity over an intron position or the end of the protein-coding region. Table S1 shows the set of species used for the molecular evolutionary analyses for each gene.

To test whether a subset of sites in a protein had evolved under recurrent positive selection, we used LRTs to compare an evolutionary model (M8) that allows a class of sites to have ω > 1 to models M7 and M8a, which allow only purifying selection or neutral evolution (Swanson et al., 2003, Yang et al., 2000). If model M8 can explain the observed sequence data significantly better than models M7 and M8a, as assessed by a LRT, then positive selection acting on a subset of sites can be inferred. For proteins for which model M8 was significantly preferred to models M7 and M8a, we used the Bayes Empirical Bayes (BEB) approach to identify at the 0.9 confidence level the specific residues that have evolved adaptively. These comparisons were done only for species within the *melanogaster* group due to the possibility of synonymous site saturation if more divergent species were included. (Such saturation occurs when the rate of substitutions at synonymous sites, *d*_S_, approaches 1, since it then becomes impossible to distinguish whether one or multiple mutations have occurred at these sites. This, in turn, reduces the accuracy of estimates of *d*_N_/*d*_S_.) To check for convergence in the ‘free-ratio’ and the sites models, we ran codeml twice with the initial omega set at 0.4 and 2, respectively.

Finally, we performed the branch-sites test for positive selection (Zhang et al., 2005) to identify classes of sites that had evolved adaptively along either of two specific lineages in the phylogeny that we identified *a priori* because they represent likely evolutionary transitions in key SP-related traits. First, we tested for sites under selection on the branch leading to the *melanogaster* group of species (Fig. 1, branch 11), since this branch corresponds with the inferred timing of when the SP receptor became expressed in the female reproductive tract and, consequently, when females became sensitive to the non-receptivity effect caused by SP (Tsuda et al., 2015). Second, we tested for sites under selection in the lineage that leads to and separates *D. ananassae* and *D. bipectinata* from the rest of the *melanogaster* group species (Fig. 1, branch 15), because these species are known to have somewhat longer sperm (Joly & Bressac, 1994, Markow, 1996, Pitnick et al., 1999). Although we inferred other important evolutionary transitions in reproductive traits on the broader *Drosophila* phylogeny (Fig. 1), we limited our branch-sites analyses to these two lineages because of the greater number of available sequenced species in the *melanogaster* group.

In the branch-sites test, we used a LRT to compare a null model allowing for only purifying and neutral selection on the focal branch with an alternative model allowing for a class of sites to evolve under positive selection (Yang, 2007, Yang & Dos Reis, 2011, Zhang et al., 2005). Recently, Venkat et al. (2018) found that this branch-sites test can have a high rate of false positives driven by multinucleotide mutations within codons (i.e., mutations at adjacent sites). To control for this issue, we implemented the tests in the Venkat model, a version of PAML developed by these authors that runs the analysis after masking these sites. PAML analyses were implemented using custom batch scripts for GNU parallel (Tange, 2018) and PAML version 4.8a or HyPhy version 2.5.1 (in the case of the Venkat model).

### Identification of seminase orthologs and paralogs

We identified the predicted amino acid sequences for orthologs of seminase, CG11037 and CG10587 in *Drosophila* species using the methods described above. To confirm that calls of orthology for seminase and its paralogs were accurate, we used Phylip’s PROML program (Felsenstein, 2005) to infer a maximum-likelihood rooted tree (using the single copies in *D. pseudoobscura* and *D. persimillis* as the outgroup, and default PROML parameters). This was consistent with the orthology assignments made using conserved gene order, except for *D. ananassae* and *D. bipectinata*, which are likely confounded by their long branch length.

### Evaluation of gene expression

*D. melanogaster, D. yakuba, D. ficusphila, D. bipectinata, D. annanassae, D. pseudoobscura* and *D. willistoni* were raised in the lab as in Tsuda et al. (2015). We CO_2_-anesthetized 9-day-old flies of each species, separated them by sex, homogenized male or female whole flies in TRIzol reagent, and purified RNA from samples and synthesized cDNA as previously described (Gubala et al., 2017). We then used species-specific primers to amplify *seminase*, *CG11037* or *CG10587*, with the non-reproductive, housekeeping gene, *ribosomal protein L32* (*RpL32*) as a control that should be expressed equally in both sexes. Genomic DNA was used as a positive control for PCR reactions, and water was used in place of template in negative control reactions.

## Results and Discussion

### SP network proteins are present in species outside of the melanogaster group

While SP orthologs have been found in species outside of the *melanogaster* group, only females of species within this group appear to show large-scale SP-mediated reproductive responses (Tsuda et al., 2015). One likely factor for this change is the evolution of *SPR* expression in the female reproductive tract in the last common ancestor of the *melanogaster* group (Tsuda et al., 2015). This evolutionary history raises the question of whether the remaining members of the SP network – all of which are critical for SP responses in *D. melanogaster* – are present outside of the group. We addressed this question bioinformatically by searching for intact orthologs across 22 *Drosophila* species with sequenced genomes.

Figure 2 shows that all SP network protein-coding gene orthologs are detectable in a large majority of the species surveyed, including those outside of the *melanogaster* group. For example, we found all currently known network proteins in *D. pseudoobscura* and *D. willistoni*, and all but one ortholog in *D. virilis*. To assess whether these orthologs were likely to function in reproduction outside of the *melanogaster* group, we examined publicly availably RNAseq data from male reproductive tracts in *D. pseudoobscura* (Yang et al., 2018) and proteomic data from male accessory glands in *D. mojavensis* (Kelleher et al., 2009). Transcripts of orthologs of male-derived network proteins were consistently enriched in (or entirely specific to) samples from whole males, male testes, and male carcasses in *D. pseudoobscura*, while showing either no or low expression in females or in male heads (Fig. S1). Expression enrichment in whole males and male carcasses is consistent with expectations for reproductive proteins produced in the male accessory gland. Some of these proteins may also be expressed in the testes, or the “testis” expression could be due to contamination of testis dissections with accessory gland tissue. The genes encoding female-derived proteins showed broader expression patterns (Fig. S1), including in whole females and whole males, but this pattern is consistent with their *D. melanogaster* orthologs, the expression of which is not limited to the female reproductive system (Brown et al., 2014). Predicted orthologs of the male-derived network proteins CG1652, CG1656, CG9997, CG17575, seminase, aquarius and antares were also identified in a proteomic analysis of the *D. mojavensis* accessory gland (Kelleher et al., 2009). Subsequent work showed that males of this species transfer transcripts of the *antares* ortholog to females during mating (Bono et al., 2011). Thus, RNAseq and proteomic data from two outgroup species are consistent with many SP network proteins functioning in reproduction outside of the *melanogaster* group.

**Figure 2.**
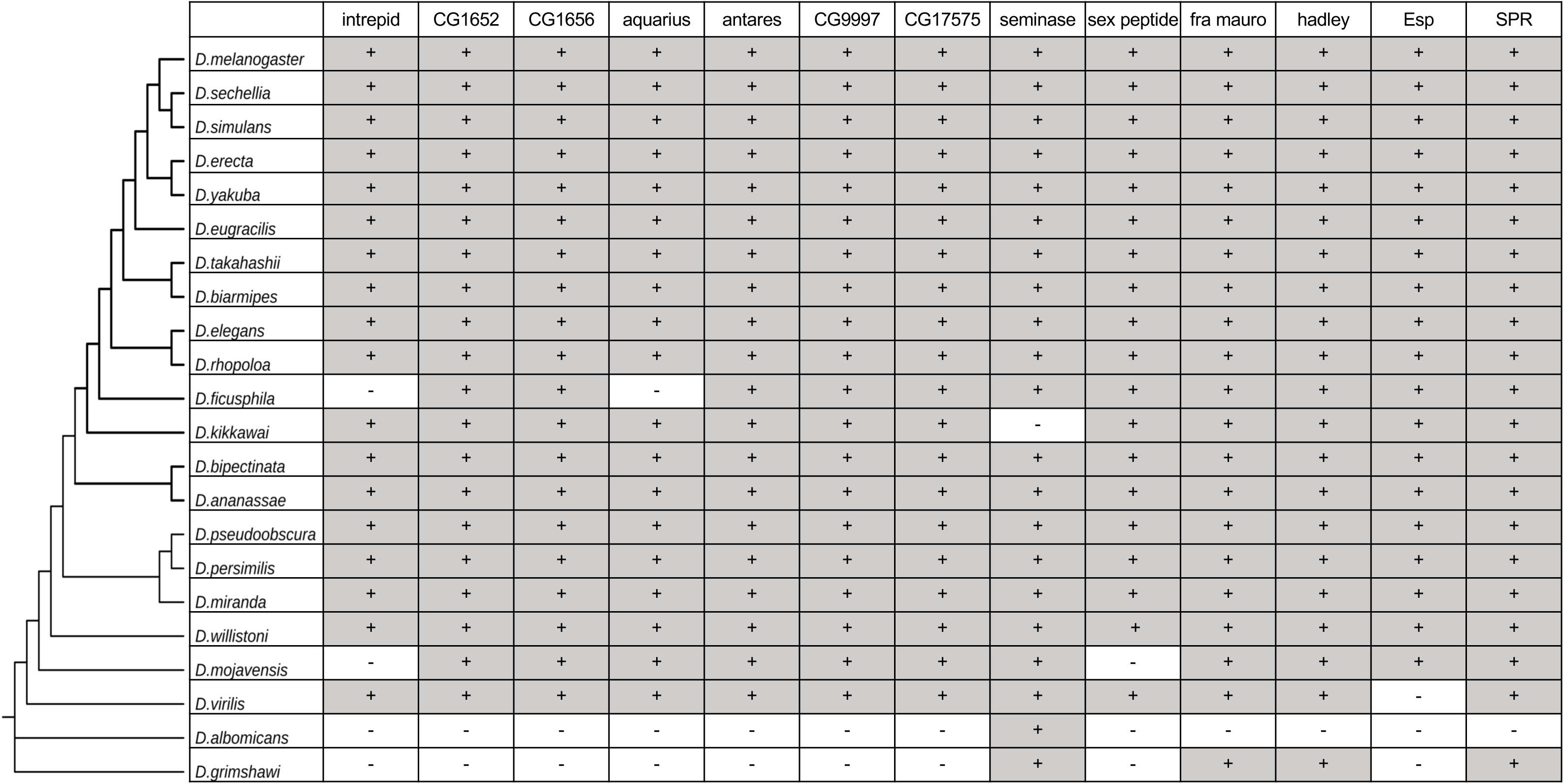
Bioinformatic identification of SP network proteins across 22 *Drosophila* species. Identified orthologs that were also reciprocal best BLAST hits are noted with a + sign, while a – sign indicates no ortholog could be identified.

It is likely that some SP network proteins function in other processes in certain species that impact their evolutionary trajectories. For example, SPR is expressed in both sexes outside of the reproductive tract (Tsuda et al., 2015, Yapici et al., 2008), and myoinhibitory peptides (MIPs) are known ligands in addition to SP (Kim et al., 2010, Poels et al., 2010, Yamanaka et al., 2010). SPR-MIP interactions outside of the reproductive tract affect sleep patterns in both sexes and remating propensity in females (Jang et al., 2017, Oh et al., 2014). Such interactions, in addition to the sexual selective pressures exerted by SP network-mediated interactions and reproductive phenotypes, have likely contributed to the evolution of SP network proteins in various *Drosophila* lineages. While most male-derived network proteins appear to have male-specific or heavily male-biased expression (in species for which expression data are available), the female-derived proteins show broader expression patterns. Understanding these proteins’ non-reproductive functions will shed additional light on evolutionary forces that may have shaped them.

### SP network proteins demonstrate evolutionary rate heterogeneity across the Drosophila *phylogeny*

Because recombination within a gene can cause false positive results in the PAML analyses, we first analyzed each set of orthologs using GARD (Kosakovsky Pond et al., 2006) to identify high-confidence recombination sites, which were detected for six of the proteins (Table S2). These six proteins were thus split into segments corresponding to the regions between recombination breakpoints, which we analyzed independently.

To begin investigating these proteins’ molecular evolution across the genus, we used PAML model M0 to estimate for each gene a single *d*_N_/*d*_S_ ratio (ω) across all species for either the full-length protein-coding sequence or for each segment identified by the GARD analysis. We then performed the branch test (Yang, 1998) to assess whether ω varied significantly across different branches of the phylogeny. Most network proteins (and segments of proteins) had full-length ω estimates around 0.2 across the full-genus tree (Table S3). While these rates are generally indicative of purifying selection, they are nonetheless faster than those observed for most proteins in *D. melanogaster* (Drosophila 12 Genomes Consortium, 2007), consistent with the general trend for reproductive proteins (Wilburn & Swanson, 2016). Three proteins showed notably slower evolutionary rates: CG17575 (most of protein found in segments with ω < 0.1), a male-expressed cysteine-rich secretory protein required for binding of SP to sperm (Ravi Ram & Wolfner, 2009); Esp (ω = 0.03), a female-expressed, predicted sulfate membrane transporter also required for long-term fertility (Findlay et al., 2014); and SPR (most of the protein contained in a segment with ω = 0.05), the female-expressed G-protein coupled receptor for SP required for female post-mating changes including egg-laying, resistance to remating and release of sperm from sperm-storage organs (Avila et al., 2015, Yapici et al., 2008). While these proteins’ slow rates of evolution could indicate that they play highly conserved roles in reproduction, it is also possible that they have evolved adaptively at only a few sites or on a few lineages (see below).

We next ran the “free ratio” model in which PAML estimates an ω value for each branch of the phylogeny. We found significant evidence of evolutionary rate heterogeneity for all but one network protein, intrepid (Table S3). Additionally, all proteins but intrepid had at least one phylogenetic branch for which ω was estimated to be > 1. While the branch test is not a rigorous test of positive selection acting on specific branches, the results indicate that the evolutionary rates of most SP network proteins have varied significantly across their evolutionary histories. Indeed, several network proteins had branch-specific ω estimates > 1 in some of the more ancestral branches of the tree, highlighting the potential for more detailed studies of these proteins’ evolutionary histories as the genomes of more divergent *Drosophila* species become available. In contrast, the constant, slow rate of evolution for intrepid implies that this protein has likely played a conserved and important role since the origin of the genus. Intrepid has undergone less functional characterization than other male-expressed male network proteins, so we cannot speculate further about its specific role(s) in reproduction.

### Several SP network proteins have undergone recurrent positive selection at specific sites since the evolution of SPR expression in female reproductive tracts

To determine the extent to which positive selection has shaped the evolution of the SP network proteins, we used the PAML sites test to ask whether any protein had a particular subset of sites that had undergone recurrent positive selection. Because of the likelihood of synonymous site saturation over longer phylogenetic distances, we limited the sequences used in this analysis to those from the *melanogaster* group. This set of species also represents the likely extent of major SP/SPR-mediated post-mating responses, as only these species express SPR at high levels in the female reproductive tract and respond to injection of synthetic SP (Tsuda et al., 2015). Thus, our analyses identify proteins that might have evolved adaptively to further improve/refine network function in the past ∼15 million years (Seetharam & Stuart, 2013).

The results of the sites analyses are shown in Table 1. Four proteins – CG9997, fra mauro, CG1652 and hadley – show significant evidence for having a class of amino acid sites that have evolved under recurrent positive selection across the *melanogaster* group of species. Three other proteins (antares, intrepid and CG17575) each have a class of sites found to be under positive selection in the Model M7/M8 comparison, but these results are no longer significant when comparing Models M8 and M8a, suggesting that the class of more quickly evolving sites identified for each protein in Model M8 may be evolving neutrally rather than under positive selection.

**Table 1.**
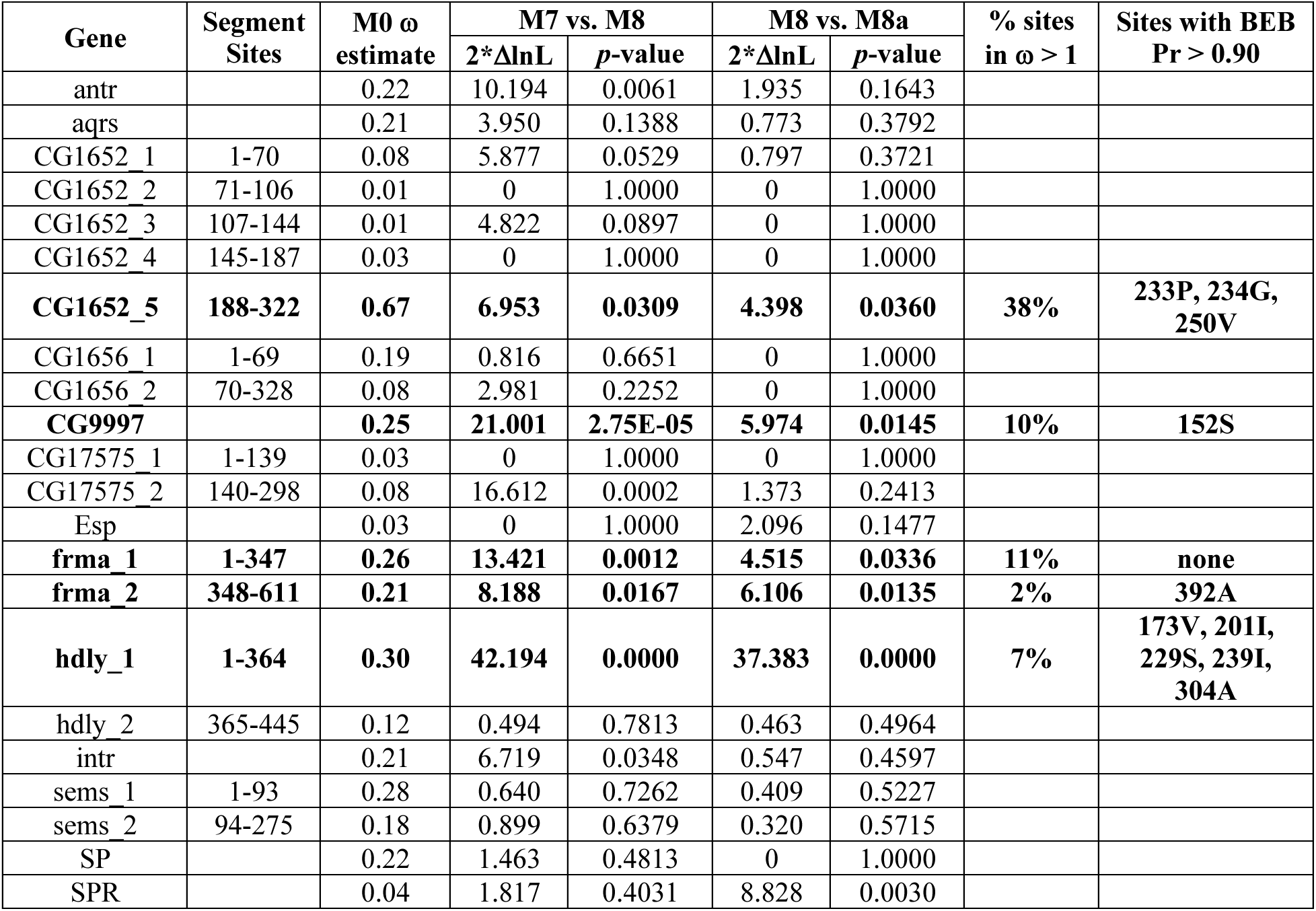
PAML sites tests for positive selection acting on SP network genes within the melanogaster group. Genes for which recombination was detected are split into numbered segments as indicated. Specific codons that were inferred to be under selection by PAML’s Bayes Empirical Bayes (BEB) analysis with Pr > 0.9 are shown for genes or segments for which positive selection was detected (i.e., in which model M8 was a significantly better fit to the data than models M7 and M8a). Amino acid site positions and identities refer to the *D. melanogaster* protein sequence.

While all members of a protein network have a degree of functional co-dependence, the male-expressed network proteins that have evolved adaptively are known to have specific, genetic interdependencies. CG9997, a serine protease homolog predicted to be catalytically inactive, must be produced in the male accessory glands for CG1652, a C-type lectin, to be transferred to mated females (Ravi Ram & Wolfner, 2009). Likewise, in the absence of CG1652, CG9997 is not efficiently “processed” from its 45-kDa form to its 36-kDa form in mated females (Ravi Ram & Wolfner, 2009, Singh et al., 2018). The loss of either protein prevents SP from accumulating on stored sperm in females. Recent work has shown that both CG9997 and CG1652 also bind to sperm, though their sperm-binding is detectable only in the hours after mating, while SP binding lasts for several days (Peng et al., 2005, Singh et al., 2018). CG9997 and CG1652 also show significant evidence of evolutionary rate covariation (Findlay et al., 2014). These results suggest that pressure to maintain their functional interactions may be a factor driving the adaptive evolution of CG9997 and CG1652, as has been observed for pairs of interacting reproductive proteins in other systems (Clark et al., 2009, Grayson, 2015).

Other work on CG9997 is consistent with its adaptive evolution. Wong et al. (2008) found evidence for recent positive selection acting on this gene by examining patterns of polymorphism and divergence between populations of *D. melanogaster* and *D. simulans*. They hypothesized that non-catalytically active serine protease homologs like CG9997 function as agonists or antagonists for active proteases. Another possibility, which is not mutually exclusive, is that protease homologs bind to other proteins or molecules in the female tract to slow their rate of digestion by active, female-derived proteases (LaFlamme & Wolfner, 2013). Under either scenario, protease homologs like CG9997 may need to continually coevolve with their interacting partners, providing the impetus for the recurrent, adaptive evolution detected here. Additionally, knockdown of *CG9997* diminishes male sperm competitive ability (Castillo & Moyle, 2014), suggesting another potential factor in its adaptive evolution.

Less functional information exists for the adaptively evolving, female-expressed proteins. Both fra mauro and hadley were identified in a screen for female-expressed proteins that coevolved with a male-expressed SP network protein; in each case, the coevolutionary signal was with CG17575 (Findlay et al., 2014). RNAi knockdown of either gene reduced female fertility, though knockdown females could receive SP and store it properly on sperm (Findlay et al., 2014). These data suggested that the proteins could be involved in maintaining the female long-term response to SP, though *fra mauro* knockdown females also showed a significant fertility defect in the 24 hrs after mating (Findlay et al., 2014). The fra mauro protein encodes a predicted neprilysin protease, which may coevolve with its as yet unknown molecular targets or antagonists (LaFlamme & Wolfner, 2013). As noted above, functional domains have not been identified for the hadley protein, so it is difficult to speculate on potential forces driving its adaptive evolution.

Notably, several proteins in the SP network showed no evidence of recurrent adaptive evolution within the *melanogaster* group, while others had subsets of sites with evolutionary rates that were elevated, but approximated neutrality. These data suggest that while some network proteins may contain regions that are under relaxed constraint, much of the functionality and interdependence of the network might have already existed at the origin of the *melanogaster* group.

### Several network proteins underwent adaptive evolution on specific lineages correlating with changes in reproductive phenotypes

While the PAML sites test described above detects recurrent adaptive evolution, protein networks can also be shaped by bursts of episodic positive selection acting on specific phylogenetic lineages. One important evolutionary transition for the SP network occurred at the base of the *melanogaster* group, when *SPR* evolved expression in the lower female reproductive tract (Tsuda et al., 2015). This change likely created (or exacerbated) a selective pressure for higher SP levels in this location, as prolonged SP-SPR signaling could promote continued egg production and prolong female non-receptivity to re-mating. Because a primary purpose of the male-expressed SP network proteins in *D. melanogaster* is to bind SP to sperm to prolong the post-mating response, we hypothesized that some of these proteins might have experienced a burst of adaptive evolution on the same phylogenetic branch on which female reproductive *SPR* expression is inferred to have evolved. Likewise, the increase in *SPR* expression in females could have created a selective pressure for other female-expressed members of the network to evolve. To test these ideas, we used the Venkat model, a modified PAML branch-sites test (Venkat et al., 2018, Zhang et al., 2005), to ask whether any network protein had a subset of sites under selection on the branch leading to the *melanogaster* group (i.e., branch 11 in Fig. 1).

Table 2 (left columns) shows the results of these tests. Two proteins show marginal evidence for adaptive evolution on branch 11, leading to the *melanogaster* group: CG1656 and SPR. As originally formulated (Zhang et al., 2005), the LRT for the branch-sites test follows a null distribution described as an equal mixture of point mass 0 and a chi-square distribution with 1 degree of freedom (df). Under this null distribution, the test statistic corresponding with a *p-* value of 0.05 is 2.71, a value exceeded in tests of both CG1656 and SPR. However, the test is typically conducted conservatively (Venkat et al., 2018, Zhang et al., 2005), following only a chi-square distribution with 1 df. The p-values listed in Table 2 are calculated based on this latter distribution, and they are marginally non-significant (0.05 < *p* < 0.1) for CG1656 and SPR.

**Table 2.** Venkat model branch-sites tests for positive selection acting on specific sites of SP network proteins on two specific lineages. P-values are calculated based on a 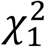 distribution. Asterisks indicate likelihood ratio test statistics that reach the *p* < 0.05 significance threshold for a null distribution derived from a 50:50 ratio of point mass 0 and the 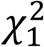distribution.

| Gene | Branch 11: to *melanogaster* group<br>(*SPR* expressed in female tract) |  |  | Branch 15: to *D. ananassae/bipictenata*<br>(increased sperm length) |  |  |
| --- | --- | --- | --- | --- | --- | --- |
| | Whole-Gene<br>$\omega$ estimate | 2*$\Delta\ln L$ | *p*-value | Whole-Gene<br>$\omega$ estimate | 2*$\Delta\ln L$ | *p*-value |
| antr | 298.77 | 2.464 | 0.116 | **6.19** | **3.948** | **0.047** |
| aqrs | 1.00 | 0 | 1.000 | 2.53 | 0.388 | 0.533 |
| CG1652 | 7.62 | 1.428 | 0.232 | 1.05 | 0.006 | 0.938 |
| CG1656 | 122.47 | 3.238* | 0.072 | 1.04 | 0 | 1.000 |
| CG9997 | 1.04 | 0 | 1.000 | 1.05 | 0.022 | 0.882 |
| CG17575 | 424.52 | 2.425 | 0.119 | 422.98 | 3.827* | 0.050 |
| Esp | 1.08 | 0 | 1.000 | 1.08 | 0 | 1.000 |
| frma | 1.08 | 0.034 | 0.854 | 1.04 | 0.012 | 0.913 |
| hdly | 1.08 | 0 | 1.000 | 1.64 | 0.008 | 0.929 |
| intr | 1.00 | 0.039 | 0.843 | 1.02 | 0 | 1.000 |
| sems | 1.00 | 0 | 1.000 | 403.51 | 0.962 | 0.327 |
| SP | 1.00 | 0 | 1.000 | 1.04 | 0.002 | 0.963 |
| SPR | 9999.99 | 3.064* | 0.080 | 1.08 | 0 | 1.000 |

The potential adaptive evolution of sites in the SPR protein along branch 11 is interesting, because this lineage also represents the time during which the protein became expressed in the female reproductive tract (Tsuda et al., 2015). Thus, it is possible that the SPR gene underwent both regulatory and protein-coding adaptations that altered how the female post-mating response is controlled. The other protein that potentially underwent adaptive evolution along this lineage is the predicted C-type lectin CG1656, which functions similarly to its recurrently rapidly evolving paralog described above, CG1652. Both lectins are required for SP’s long-term binding to stored sperm, and both proteins themselves bind sperm temporarily in the hours after mating (Singh et al., 2018). Given the potential selective pressure to bind more SP to stored sperm in female tracts expressing *SPR*, it is possible that the adaptive evolution of CG1656 on this key phylogenetic branch could have helped to improve the efficiency of SP’s binding to sperm. This idea could be tested in future experiments by either identifying and mutating the residues likely to have changed along branch 11 and/or by substituting an outgroup ortholog of CG1656 (and potentially its duplicate, CG1652) into *D. melanogaster* and examining the effects on SP’s sperm binding and on the female long-term post-mating response.

Prior work demonstrated that SP binds to the full length of *D. melanogaster* sperm (Peng et al., 2005, Ravi Ram & Wolfner, 2009, Singh et al., 2018). Indeed, the ability of SP (and potentially other molecules) to bind sperm and then influence post-mating responses is one hypothesis for why sperm tails have evolved to be so long in many *Drosophila* species. Within the *melanogaster* group species that experience SP-mediated post-mating responses, one notable change in reproductive physiology is that the sperm of *D. ananassae* and its closely related species are considerably longer than those of *D. melanogaster* (*D. ananassae* sperm length: 3.3 mm; *D. melanogaster* and other *melanogaster* group species sperm length: just under 2 mm (Pitnick et al., 1999, Joly & Bressac, 1994, Markow, 1996)). We thus infer that a major (>50%) increase in sperm length occurred on the branch of the phylogeny leading to *D. ananassae* and *D. bipectinata* (branch 15 in Fig. 1).

To test for whether any SP network proteins experienced adaptive evolution concurrent with this change in sperm length, we again used the modified branch-sites test. Two network proteins, antares and CG17575, show evidence of positive selection acting on specific sites on the lineage leading to *D. ananassae* and *D. bipectinata* (Table 2, right columns). Antares’ signal of selection is significant under both null distributions described above, while CG17575’s signal is significant under the mixed null distribution and approached significance (*p* = 0.0504) under the conservative test. In addition to facilitating SP’s long-term binding to sperm, antares also binds to sperm itself for a shorter period (Findlay et al., 2014, Singh et al., 2018). Thus, antares might have evolved adaptively to facilitate greater or more efficient binding of either itself or SP to sperm as sperm tails lengthened. Interestingly, the antares ortholog in outgroup species *D. mojavensis* and *D. arizonae* was also found to evolve under diversifying selection (Bono et al., 2015), even though *D. mojavensis* does not have a currently detectable SP ortholog (Tsuda et al., 2015) (Fig. 2). Heterospecific matings between these species fail due to post-mating, pre-zygotic isolating barriers, which include problems with sperm storage in the female reproductive tract (Kelleher & Markow, 2007). It is thus possible that antares plays an essential role in binding molecules to sperm and/or facilitating sperm storage, and that the male reproductive activity of antares has been refined by different selective pressures in different lineages.

CG17575 is a male-expressed, cysteine-rich secretory protein required for SP and other sperm-binding network proteins to localize from the female uterus, where seminal proteins and sperm are first deposited, into the seminal receptacle (SR), the primary site of sperm storage in *D. melanogaster* (Ravi Ram & Wolfner, 2009, Singh et al., 2018). Since CG17575 does not itself bind sperm (Singh et al., 2018), further details of how CG17575 provides for proper localization of other seminal proteins to the seminal receptacle are needed before we can speculate on the selective forces that might have contributed to its evolution in this lineage.

The branch-sites tests for branches 11 and 15 reported above were conducted using full-length gene sequences, since the test has limited power. However, we repeated this analysis on all segments of the six genes for which recombination was detected. These results (Table S4) found marginal evidence for selection for antares on branch 11 and for a segment of CG1652 on branch 15. CG1656 was not among the genes for which recombination was detected (Table S2), so its results above are unaltered.

### Seminase gene duplicates retain male-specific expression patterns across melanogaster group species

In addition to CG17575, the male-expressed serine protease seminase is required for the localization of SP and other male-expressed proteins to the SR after mating (LaFlamme et al., 2012, Singh et al., 2018). Seminase arose through gene duplication in the lineage leading to the *melanogaster* group of flies. The genomes of *D. pseudoobscura* and other outgroup species contain only one detectable copy of the gene, but in *D. melanogaster* and its fellow *melanogaster* group members, there are three tandemly arrayed, intron-containing copies, suggesting two distinct DNA-based duplication events. The other genes are *CG10587* and *CG11037*. Like *seminase*, both are expressed specifically in the male accessory gland in *D. melanogaster* (Brown et al., 2014, Leader et al., 2018). While we detected no recurrent or episodic positive selection acting on seminase after these duplications (Tables 1-2), we were curious whether it or its paralogs might have evolved different expression patterns (and, thus, potential functions) after duplication. We thus performed RT-PCR to amplify each paralog from cDNA isolated from males or females of a variety of species from the *melanogaster* group. We also assessed the expression of the single-copy parent gene from *D. pseudoobscura* and *D. willistoni*. Our results (Figure 3) show that both the single-copy genes from the outgroup species, as well as all of paralogs from all *melanogaster* group species tested, are expressed specifically in adult males. This result is consistent with the ancestral single copy of seminase also functioning in male reproduction (and potentially with other SP network proteins).

**Figure 3.**
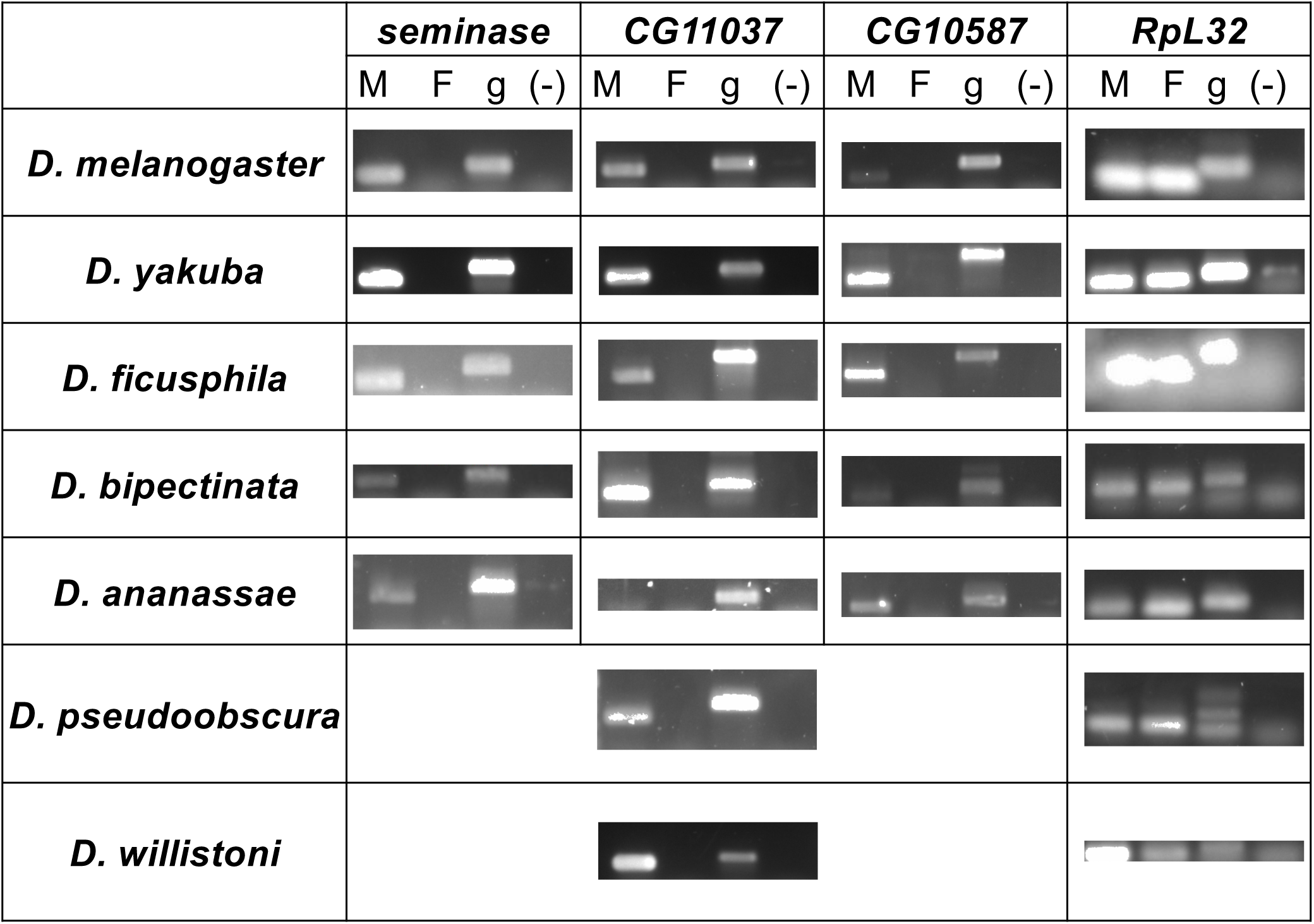
RT-PCR on *seminase* and paralogs shows conserved, male-biased expression after duplication. Orthologs of *seminase*, *CG10586* and *CG11037* show male-specific expression in various *melanogaster* group species, though the level of expression between paralogs and species is somewhat variable. The single-copy parent gene in *D. pseudoobscura* and *D. willistoni* is also expressed in a male-specific manner. The non-reproductive housekeeping gene, *RpL32*, was included as a positive control for successful cDNA synthesis from both sexes.

Given that seminase itself has additional reproductive functions beyond its role in the SP network (LaFlamme et al., 2012), it is possible that the paralogs have sub- or neo-functionalized to have unique roles, in spite of their conserved expression patterns. Future studies should evaluate how the paralogs contribute to reproduction, which may suggest possible evolutionary forces that affected their evolution after the gene duplication events.

## Conclusions

Sex peptide is directly responsible for major changes in female post-mating behavior and physiology and is therefore one of the best characterized reproductive proteins to date. SP-mediated responses appear to have arisen specifically in the *melanogaster* group of *Drosophila*, and they manifest in full only with the help of a suite of male- and female-derived proteins, the SP network. We have shown that these proteins are present and expressed in species outside of the *melanogaster* group, suggesting they likely function in reproduction in these species and that they did so in a common ancestor. Within the *melanogaster* group, several network proteins (CG9997, CG1652, fra mauro, and hadley) have experienced recurrent positive selection, suggesting that continued, adaptive evolution refined SP network function. A non-overlapping set of proteins, including CG1656, SPR, antares, and CG17575, showed some evidence of bursts of adaptive evolution on specific phylogenetic lineages corresponding with major changes in SP network reproductive phenotypes. Taken together, these data suggest that SP network proteins may have interacted to affect reproduction before the evolution of major SP-mediated changes in the *melanogaster* group. However, once SPR became expressed at high levels in the female reproductive tract in the common ancestor of this group (Tsuda et al., 2015), a combination of both quick bursts of adaptation on specific lineages and recurrent changes at specific protein sites helped the network evolve into the present form observed in *D. melanogaster*. This study demonstrates how changes in both regulatory and protein-coding regions can affect the evolution of protein networks and motivates future functional studies of the SP network proteins in *Drosophila* species both within and outside of the *melanogaster* group.

## Acknowledgements

We thank L. Moyle, M. Hahn, A. Venkat and M. Wu for assistance in implementing the Venkat model, D. Thakral for help developing batch PAML scripts, K. Ober and M. Wolfner for helpful discussions, and two reviewers for valuable feedback on the manuscript. This work was supported by NSF CAREER Award 1652013 and a summer research fellowship generously provided by Dr. William Crowley. The authors report no conflicts of interest.

## Supplemental Materials

**Table S1. Orthologs used for each gene in PAML analyses**. Some orthologs that were identified in Table 2 were excluded from PAML analysis due to unresolved intron borders and/or poor alignment quality. Only species above the dotted line (the *melanogaster* group) were analyzed in the sites tests.

**Table S2. GARD results showing inferred recombination breakpoints.** Breakpoint positions refer to nucleotide positions in the alignment files used. However, since alignments include gaps, these positions do not necessarily have a 3:1 correspondence with the *D. melanogaster* amino acid positions reported in Tables 1-3. The first table shows recombination breakpoints detected for aligned sequences from the entire *Drosophila* genus, which were used for the branch and branch-sites tests. The second table shows recombination breakpoints detected for aligned sequences from only the *melanogaster* group, which were used for the sites tests.

**Figure S1. RNAseq data from D. pseudoobscura support reproductive functions for SP network proteins in a species that lacks full-scale SP responses.** A) *D. pseudoobscura* expression patterns for each member of the SP network. Dark shading indicates high expression levels, stripes indicate low (but detectable) expression, and no shading indicates no expression detected in a given sample. Male-derived network proteins show male-biased or male-specific expression, consistent with reproductive functions. B) Examples of *D. pseudoobscura* expression data for several SP network genes; shading in part (A) is based on these data. The RNAseq data were accessed via FlyBase and generated by Yang et al. (Yang et al., 2018).

**Table S3. Branch tests for rate heterogeneity.** Partitions were implemented in PAML analyses if they were significant in both the LH and RH tests.

**Table S4. Venkat model branch-sites tests for positive selection acting on specific sites of SP network proteins detected by GARD to have multiple recombination segments.** The table shows results for both branch 11 and branch 15 tests. Only the six genes for which recombination was detected in the relevant trees are shown in the table. *P-*values are calculated based on a 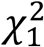 distribution. Asterisks indicate likelihood ratio test statistics that reach the *p* < 0.05 significance threshold for a null distribution derived from a 50:50 ratio of point mass 0 and the 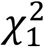 distribution.

