## Supplemental material for "Molecular Evolution of the Sex Peptide Network in *Drosophila*"

**Supplemental Materials**

**Table S1. Orthologs used for each gene in PAML analyses**. Some orthologs that were identified in Table 2 were excluded from PAML analysis due to unresolved intron borders and/or poor alignment quality. Only species above the dotted line (the *melanogaster* group) were analyzed in the sites tests.

**Table S2. GARD results showing inferred recombination breakpoints.** Breakpoint positions refer to nucleotide positions in the alignment files used. However, since alignments include gaps, these positions do not necessarily have a 3:1 correspondence with the *D. melanogaster* amino acid positions reported in Tables 1-3. The first table shows recombination breakpoints detected for aligned sequences from the entire *Drosophila* genus, which were used for the branch and branch-sites tests. The second table shows recombination breakpoints detected for aligned sequences from only the *melanogaster* group, which were used for the sites tests.

**Table S3. Branch tests for rate heterogeneity.** Partitions were implemented in PAML analyses if they were significant in both the LH and RH tests.

**Table S4. Venkat model branch-sites tests for positive selection acting on specific sites of SP network proteins detected by GARD to have multiple recombination segments.** The table shows results for both branch 11 and branch 15 tests. Only the six genes for which recombination was detected in the relevant trees are shown in the table. *P*-values are calculated based on a $\chi_1^2$ distribution. Asterisks indicate likelihood ratio test statistics that reach the $p < 0.05$ significance threshold for a null distribution derived from a 50:50 ratio of point mass 0 and the $\chi_1^2$ distribution.

**Figure S1. RNAseq data from *D. pseudoobscura* support reproductive functions for SP network proteins in a species that lacks full-scale SP responses.** A) *D. pseudoobscura* expression patterns for each member of the SP network. Dark shading indicates high expression levels, stripes indicate low (but detectable) expression, and no shading indicates no expression detected in a given sample. Male-derived network proteins show male-biased or male-specific expression, consistent with reproductive functions. B) Examples of *D. pseudoobscura* expression data for several SP network genes; shading in part (A) is based on these data. The RNAseq data were accessed via FlyBase and generated by Yang et al. (Yang et al., 2018).

**Table S1**

| | intrepid | CG1652 | CG1656 | aquarius | antares | CG9997 | CG17575 | seminase | SP | fra mauro | hadley | Esp | SPR |
|---|---|---|---|---|---|---|---|---|---|---|---|---|---|
| *D. melanogaster* | X | X | X | X | X | X | X | X | X | X | X | X | X |
| *D. sechellia* | X | X | X | X | X | X | X | X | X | X | X | X | X |
| *D. simulans* | X | X | X | X | X | X | X | X | X | X | X | X | X |
| *D. erecta* | X | X | X | X | X | X | X | X | X | X | X | X | X |
| *D. yakuba* | X | X | X | X | X | X | X | X | X | X | X | X | X |
| *D. eugracilis* | X | X | X | X | X | X | X | X | X | X | | | X |
| *D. takahashii* | X | X | X | X | X | X | X | X | X | | X | X | X |
| *D. biarmipes* | X | X | X | X | X | X | X | X | X | X | X | X | X |
| *D. elegans* | X | X | X | X | X | X | X | X | X | X | | | X |
| *D. rhopoloa* | X | X | X | X | X | X | X | X | X | X | X | X | X |
| *D. ficusphila* | | X | X | | X | | X | X | X | X | | X | X |
| *D. kikkawai* | X | X | X | X | | X | X | | X | X | | | X |
| *D. bipectinata* | X | X | X | X | X | X | X | X | X | X | | X | X |
| *D. ananassae* | X | X | X | X | X | X | X | X | X | X | X | X | X |
| *D. pseudoobscura* | X | X | X | X | X | X | X | X | X | X | X | X | X |
| *D. persimilis* | X | X | X | X | X | X | X | X | X | X | X | X | |
| *D. miranda* | X | X | X | X | | X | X | X | X | X | | X | X |
| *D. willistoni* | X | X | X | X | X | X | X | X | X | X | X | X | X |
| *D. mojavensis* | | X | X | X | | X | X | X | | X | X | X | X |
| *D. virilis* | | X | X | X | X | X | X | X | X | X | X | | X |
| *D. albomicans* | | | | | | | | X | | | | | |
| *D. grimshawi* | | | | | | | | X | | X | X | | X |

**Table S2.** Recombination breakpoints detected by GARD.

| Branch & Branch-Sites Tree Recombination | | | | | |
|---|---|---|---|---|---|
| **Gene** | **AICc** | **ΔAICc** | **Breakpoint** | **LH p-value** | **RH p-value** |
| ***antr*** | **20700.5** | **51.118** | 184 | 0.1088 | 0.0512 |
| | | | **309** | **0.0100** | **0.0132** |
| ***aqrs*** | **29282.2** | **35.937** | **83** | **0.0034** | **0.0068** |
| ***CG1652*** | **26798.8** | **169.498** | 631 | 1.0000 | 0.0004 |
| | | | **1309** | **0.0376** | **0.0004** |
| *CG1656* | 30121.1 | 57.130 | 327 | 0.0008 | 0.1720 |
| | | | 516 | 0.3144 | 0.0001 |
| | | | 720 | 1.0000 | 0.0008 |
| | | | 1239 | 1.0000 | 0.0008 |
| *CG9997* | 25425.1 | 116.194 | 164 | 0.6920 | 0.0004 |
| | | | 280 | 0.0004 | 1.0000 |
| ***CG17575*** | **14623.8** | **58.081** | **192** | **0.0112** | **0.0004** |
| | | | **435** | **0.0016** | **0.0004** |
| *Esp* | 26251.0 | 13.370 | 366 | 0.4656 | 0.3096 |
| | | | 744 | 0.0312 | 1.0000 |
| | | | 1017 | 0.6108 | 0.0120 |
| | | | 1200 | 0.1596 | 0.0048 |
| | | | 1386 | 0.0012 | 1.0000 |
| | | | 1863 | 0.1788 | 0.0048 |
| *frma* | 57439.4 | 51.270 | 1100 | 0.0600 | 0.3280 |
| | | | 1471 | 0.6312 | 0.0424 |
| | | | 1752 | 1.0000 | 0.1088 |
| | | | 1908 | 0.5168 | 1.0000 |
| ***hdly*** | **26370.8** | **51.700** | 162 | 0.0960 | 0.0030 |
| | | | 275 | 0.0010 | 0.0800 |
| | | | 574 | 1.0000 | 1.0000 |
| | | | 1344 | 0.6100 | 0.0030 |
| | | | **1464** | **0.0280** | **0.0010** |
| *intr* | 18076.1 | 75.453 | 217 | 0.0002 | 0.2408 |
| *sems* | 27192.1 | 31.359 | 198 | 0.0002 | 0.1684 |
| *SP* | *no recombination sites detected* | | | | |
| ***SPR*** | **24964.3** | **61.371** | **187** | **0.0002** | **0.0124** |

### Table S2, cont.

| Sites Tree Recombination | | | | | |
|---|---|---|---|---|---|
| **Gene** | **AICc** | **ΔAICc** | **Breakpoint** | **LH p-value** | **RH p-value** |
| *antr* | 10610.20 | 9.47 | 198 | 0.4648 | 0.3812 |
| | | | 381 | 1.0000 | 1.0000 |
| *aqrs* | 21417.90 | 20.30 | 171 | 0.5144 | 0.0004 |
| | | | 263 | 0.0004 | 0.4032 |
| ***CG1652*** | **14821.77** | **8.14** | **232** | **0.0152** | **0.0216** |
| | | | **341** | **0.0008** | **0.0096** |
| | | | **456** | **0.0008** | **0.0048** |
| | | | **587** | **0.0120** | **0.0008** |
| ***CG1656*** | **18109.10** | **20.85** | 180 | 0.1632 | 0.0006 |
| | | | **243** | **0.0006** | **0.0144** |
| | | | 429 | 0.0138 | 0.2610 |
| *CG9997* | 16730.20 | 149.98 | 163 | 0.6864 | 0.0004 |
| | | | 295 | 0.0004 | 0.5576 |
| ***CG17575*** | 10005.10 | 12.69 | 315 | 0.1732 | 0.0084 |
| | | | **432** | **0.0176** | **0.0004** |
| ***frma*** | **36579.94** | **5.60** | 434 | 1.0000 | 1.0000 |
| | | | 1020 | 0.4480 | 0.0168 |
| | | | 1188 | 0.4704 | 0.0014 |
| | | | **1251** | **0.0014** | **0.0098** |
| | | | 1419 | 0.1022 | 0.1008 |
| | | | 1535 | 1.0000 | 1.0000 |
| | | | 1663 | 0.0070 | 1.0000 |
| ***hdly*** | **13041.84** | **9.69** | 150 | 0.1704 | 0.4956 |
| | | | 268 | 0.1584 | 1.0000 |
| | | | 355 | 1.0000 | 1.0000 |
| | | | 540 | 1.0000 | 1.0000 |
| | | | 1054 | 0.6408 | 0.1572 |
| | | | **1201** | **0.0012** | **0.0012** |
| *intr* | 14104.50 | 89.18 | 202 | 0.0002 | 0.8628 |
| ***sems*** | **15379.20** | **8.29** | 135 | 0.0004 | 1.0000 |
| | | | **288** | **0.0036** | **0.0002** |
| *SP* | 3627.47 | 34.40 | 146 | 0.0284 | 0.1888 |
| *SPR* | 14968.20 | 21.27 | 197 | 1.0000 | 0.0524 |
| | | | 294 | 0.0004 | 1.0000 |

**Table S3.** Branch tests for rate heterogeneity.

| Gene | | Whole gene $\omega$ | "Free ratio" model | | | |
|---|---|---|---|---|---|---|
| | | | $2*\Delta\ln L$ | df | p-value | Branches with $\omega > 1$ |
| ***antr_1*** | 1-74 | 0.22 | 60.522 | 31 | 0.0011 | 7, 11, 17, 18, 20, 25, 34 |
| ***antr_2*** | 75-272 | 0.17 | 49.802 | 31 | 0.017 | 7, 8, 11, 17, 21, 24, 26, 25, 33, 37, 40 |
| ***aqrs_1*** | 1-150 | 0.21 | 54.648 | 35 | 0.018 | 5, 10, 13, 14 |
| ***aqrs_2*** | 151-366 | 0.19 | 59.303 | 35 | 0.006 | 5, 8, 12, 14 |
| ***CG1652_1*** | 1-298 | 0.21 | 82.144 | 37 | 2.384E-05 | 5, 13 |
| *CG1652_2* | 299-322 | 0.62 | 48.614 | 37 | 0.095 | |
| ***CG1656*** | | 0.30 | 93.766 | 36 | 4.82E-07 | 5, 7, 13, 23 |
| ***CG9997*** | | 0.25 | 76.726 | 35 | 5.92E-05 | 5, 12, 34, 36 |
| *CG17575_1* | 1-54 | 0.14 | 38.267 | 37 | 0.411 | |
| ***CG17575_2*** | 55-135 | 0.01 | 60.917 | 37 | 0.008 | 4, 5, 12, 14, 16, 19, 21, 24, 25, 34, 35, 38 |
| ***CG17575_3*** | 136-298 | 0.08 | 61.803 | 37 | 0.006 | 5, 8 |
| ***Esp*** | | 0.03 | 77.571 | 29 | 2.61E-06 | 7 |
| ***frma*** | | 0.23 | 185.074 | 37 | 1.37E-21 | 3, 5, 11, 12, 14, 23 |
| ***hdly_1*** | 1-362 | 0.24 | 77.501 | 27 | 8.97E-07 | 3, 5, 17, 29 |
| ***hdly_2*** | 363-445 | 0.01 | 41.731 | 27 | 0.0349 | 3, 5, 7, 21, 25, 27, 29, 35, 38 |
| *intr* | | 0.21 | 43.155 | 30 | 0.057 | |
| ***sems*** | | 0.20 | 79.122 | 39 | 1.53E-04 | 2, 7, 10, 15, 38 |
| ***SP*** | | 0.14 | 73.678 | 35 | 1.43E-04 | 13, 27, 33, 35 |
| *SPR_1* | 1-6 | 0.20 | 28.051 | 37 | 0.855 | |
| ***SPR_2*** | 7-429 | 0.05 | 177.989 | 37 | 2.41E-20 | 1, 11, 15, 20, 24, 27 |

Branches 1,3,5,7 = ancestors of mel group and others; branches 2,4,6,8,10,12,13,16,17 = leading to non-*mel* group species; all other branches: origin of (11) or within *mel* group.

**Table S4**. Venkat model branch-sites tests for positive selection acting on specific sites of SP network proteins detected by GARD to have multiple recombination segments.

| Branch 11: to *melanogaster* group | | | | |
|---|---|---|---|---|
| Gene | Sites | Segment ω | Venkat Model | |
| | | | 2*ΔlnL | $\chi_1^2$ p-value |
| *antr_1* | 1-74 | 1.083 | 0 | 1.0000 |
| *antr_2* | 75-272 | 12.323 | 3.119* | 0.0774 |
| *aqrs_1* | 1-150 | 1.043 | 0.002 | 0.9609 |
| *aqrs_2* | 151-366 | 1.041 | 0.002 | 0.9643 |
| *CG1652_1* | 1-298 | 1.002 | 0.008 | 1.0000 |
| *CG1652_2* | 299-322 | 1.000 | 0.005 | 0.9459 |
| *CG17575_1* | 1-54 | 1.086 | 0 | 1.0000 |
| *CG17575_2* | 55-135 | 1.083 | 0.003 | 0.9549 |
| *CG17575_3* | 136-298 | 264.710 | 1.435 | 0.2310 |
| *hdly_1* | 1-362 | 1.041 | 0.002 | 0.9643 |
| *hdly_2* | 363-445 | 1.088 | 0 | 1.0000 |
| *SPR_1* | 1-6 | 12.83 | 0 | 1.0000 |
| *SPR_2* | 7-429 | 1.083 | 0.013 | 0.9106 |

| Branch 15: to *D. ananassae/bipectinata* | | | | |
|---|---|---|---|---|
| Gene | Sites | Segment ω | Venkat Model | |
| | | | 2*ΔlnL | $\chi_1^2$ p-value |
| *antr_1* | 1-74 | 0.0755 | 0.002 | 0.9681 |
| *antr_2* | 75-272 | 1.086 | 0 | 1.0000 |
| *aqrs_1* | 1-150 | 0.0138 | 0 | 1.0000 |
| *aqrs_2* | 151-366 | 0.0348 | 0.094 | 0.7594 |
| *CG1652_1* | 1-298 | 0.6712 | 2.762* | 0.0965 |
| *CG1652_2* | 299-322 | 0.1889 | 0 | 1.0000 |
| *CG17575_1* | 1-54 | 0.0817 | 1.392 | 0.2380 |
| *CG17575_2* | 55-135 | 0.0317 | 0.001 | 0.9724 |
| *CG17575_3* | 136-298 | 0.0787 | 0 | 1.0000 |
| *hdly_1* | 1-362 | 0.3000 | 0.000 | 1.0000 |
| *hdly_2* | 363-445 | 0.1194 | 0.117 | 0.7325 |
| *SPR_1* | 1-6 | 7.115 | 0 | 1.0000 |
| *SPR_2* | 7-429 | 1.000 | 0.005 | 0.9436 |

**Fig S1A**

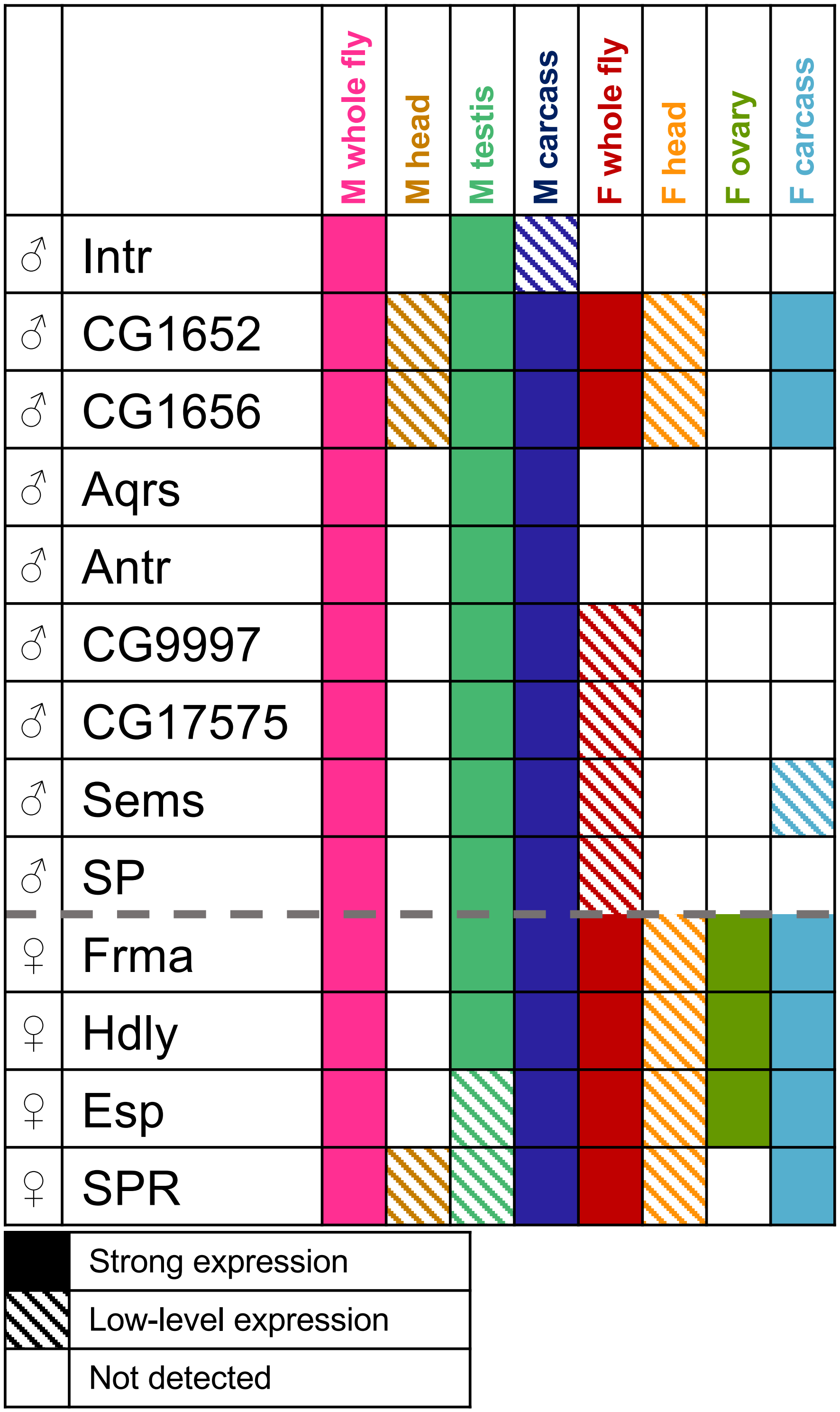
M whole fly
M head
M testis
M carcass
F whole fly
F head
F ovary
F carcass
♂ Intr
♂ CG1652
♂ CG1656
♂ Aqrs
♂ Antr
♂ CG9997
♂ CG17575
♂ Sems
♂ SP
♀ Frma
♀ Hdly
♀ Esp
♀ SPR
Strong expression
Low-level expression
Not detected

### Fig S1B

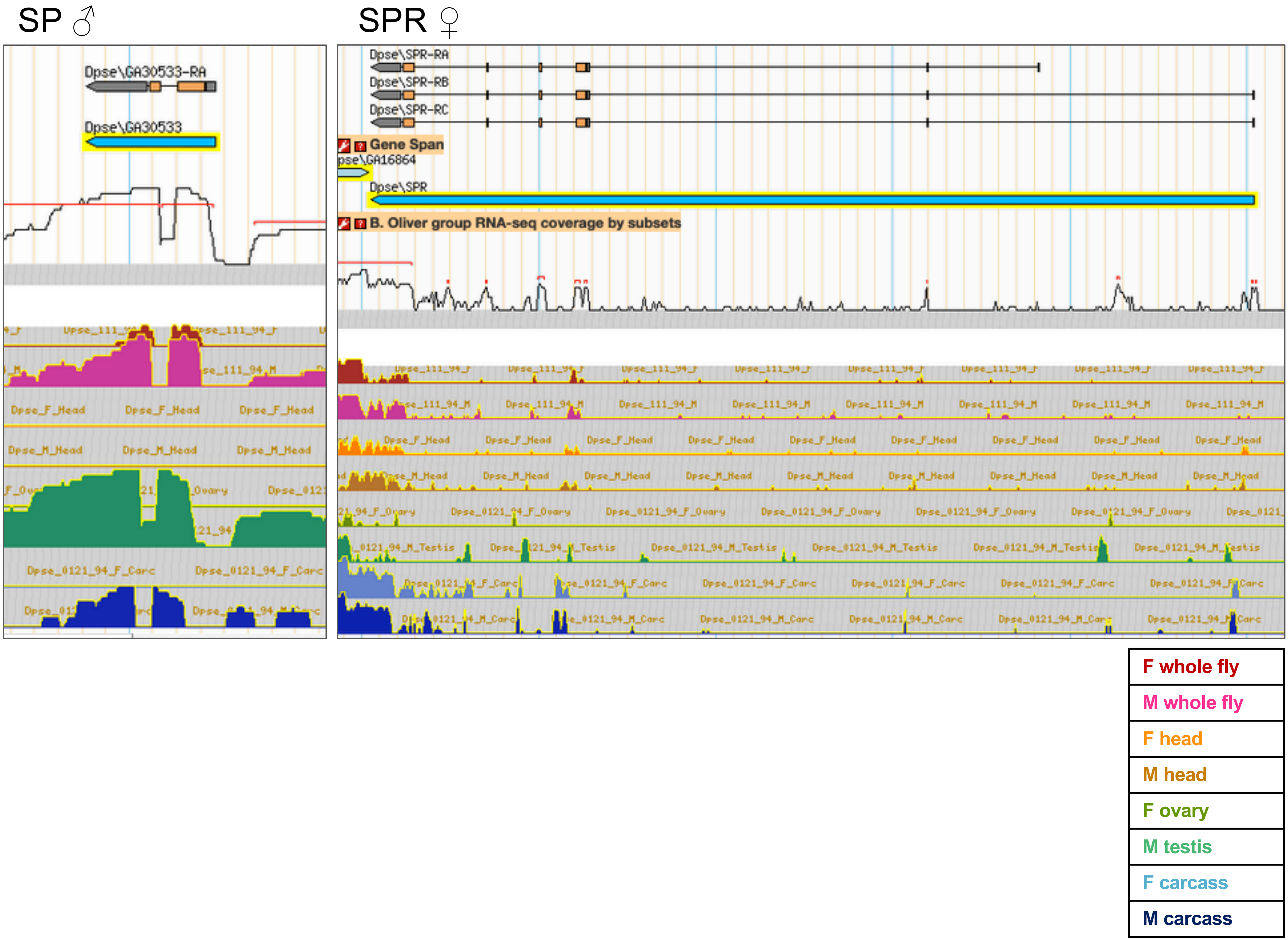

SP ♂
SPR ♀
Dpse\GA30533-RA
Dpse\GA30533
Dpse\SPR-RA
Dpse\SPR-RB
Dpse\SPR-RC
Gene Span
pse\GA16864
Dpse\SPR
B. Oliver group RNA-seq coverage by subsets
Dpse_F_Head
Dpse_M_Head
Dpse_0121_94_F_Carc
Dpse_111_94_F
Dpse_111_94_M
Dpse_0121_94_F_Ovary
Dpse_0121_94_M_Testis
Dpse_0121_94_M_Carc
F whole fly
M whole fly
F head
M head
F ovary
M testis
F carcass
M carcass

### Fig S1B, cont.

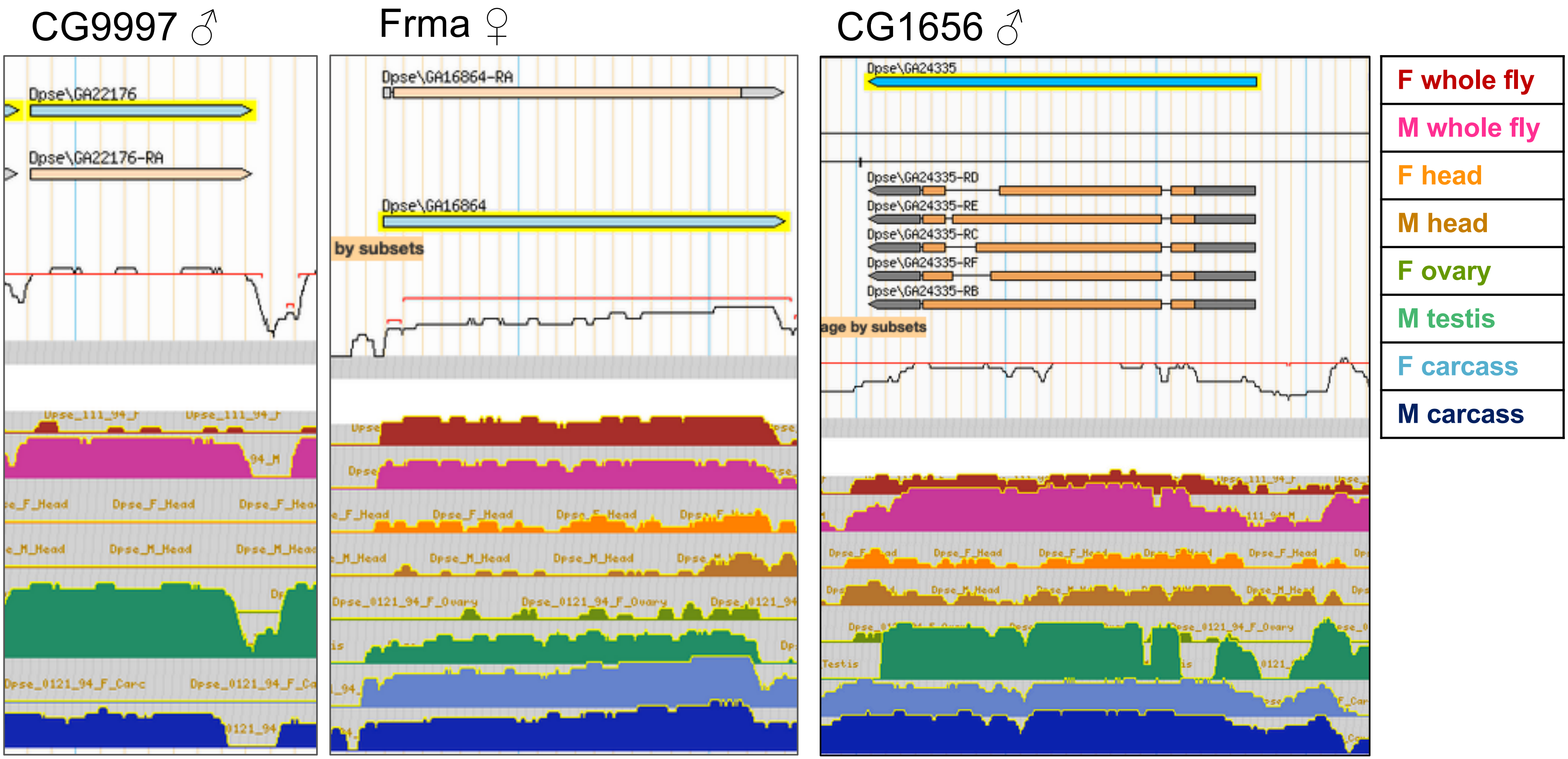
CG9997 ♂
Frma ♀
CG1656 ♂
Dpse\GA22176
Dpse\GA22176-RA
Dpse\GA16864-RA
Dpse\GA16864
by subsets
Dpse\GA24335
Dpse\GA24335-RD
Dpse\GA24335-RE
Dpse\GA24335-RC
Dpse\GA24335-RF
Dpse\GA24335-RB
F whole fly
M whole fly
F head
M head
F ovary
M testis
F carcass
M carcass
